# Isolation and Characterization of Potential Pathogenic Fungi Associated with Watermelons in Southwest Mississippi

**DOI:** 10.1101/2025.09.05.674577

**Authors:** Zharia Barnes, Prachi Bista, Glennescia Tenner, Lewis Brooks, Frank Mrema, Bed Prakash Bhatta

## Abstract

Watermelon is an important specialty crop in the USA. Fungal diseases primarily challenge the production and productivity of watermelon. The understanding of prevailing diseases, gained through genotypic and phenotypic techniques, eventually helps researchers develop disease management strategies. This study involved scouting and sampling of a watermelon research field in Southwest Mississippi for diseases. Nine distinct potential pathogenic fungi were identified from symptomatic leaves and fruits through development of pathogen pure cultures followed by PCR amplification of internal transcribed spacer regions and sequencing. The identified fungi were *Didymella americana, Diaporthe melonis, Fusarium incarnatum, Alternaria longissima, Curvularia pseudobrachyspora, Xylaria multiplex, Phoma* sp., *Nigrospora sphaerica*, and *Fusarium chlamydosporum*. The fungal isolates were ascomycetes at the phylum-level and belonged to the orders of fungi: Pleosporales, Hypocreales, Diaporthales, Xylariales, and Trichosphaeriales. Future research studies include pathogenicity assays with a broader host range, whole genome sequencing to develop deeper understanding of the pathogenic fungi, and formulation of disease management strategies for high-yielding, superior quality, and profitable watermelon production.

## Introduction

Watermelons originated in northeast or west Africa and are now cultivated globally [1]. The domesticated, edible and sweet dessert watermelon is referred to as *Citrullus lanatus* [2]. The wild progenitor of watermelons is *C. mucosospermus* or ‘egusi’ watermelon which is also known for its edible seeds [1]. The bitter-flavored wild watermelons are part of *Citrullus amarus* or ‘citron’ melon and *Citrullus colocynthis* [3]. Fungal diseases have been historically associated with a decline in production, productivity, and quality of watermelons and are the major biotic limiting factors affecting watermelon production [4-6]. Fungal pathogens can overwinter on infected debris and persist in the soil for many years [7]. Though fungicide application may be an effective management strategy to curb the losses with fungal diseases, crop improvement via disease resistance breeding remains the safer, environmentally friendly, cost-effective, and sustainable approach of disease management [8]. Isolation, identification, and characterization of fungal pathogens of watermelon and associated cucurbits help to improve disease resistance, yield, and produce quality.

The major fungal diseases affecting the production and marketability of watermelons in the U.S. are Fusarium wilt caused by *Fusarium oxysporum* f. sp. *niveum*, gummy stem blight caused by *Didymella bryoniae*, powdery mildew caused by *Podosphaera xanthii*, and anthracnose caused by *Colletotrichum orbiculare* [9-17]. The incidence, severity, and subsequent yield losses occurring due to fungal diseases are devastating. For instance, the disease severity of anthracnose in research plots in Georgia ranged more than 90% [18]. Another example is the ability of Fusarium wilt pathogen’s ability to thrive in the soil for years in the form of chlamydospores and develop resistance against fungicides [19]. Besides these, many diseases on cucurbits caused by novel fungal species have emerged in recent years. These include leaf spot caused by *Curvularia pseudobrachyspora* [20] and fruit rot and stem canker caused by *Diaporthe* species [21].

Identification of fungi using molecular techniques is critical for their characterization. The internal transcribed spacer (ITS) is a conserved, non-coding region of DNA located between the small subunit (18S) and the large sub-unit (28S) of fungal ribosomal RNA (rRNA). The ITS region has been proposed as a primary fungal barcode marker because of its ability to successfully identify a broad range of fungi [22]. The fungal primers ITS-5 and ITS-4 are two of the primers located in this region besides ITS-1, ITS-2, and ITS-3 [23]. The primers in the ITS region can amplify the DNA template of both ascomycetes and basidiomycetes [24]. The positions of the two primers, ITS-5 and ITS-4 on fungal ribosomal RNA (rRNA) or rDNA is shown in Figure 1.

**Figure 1.**
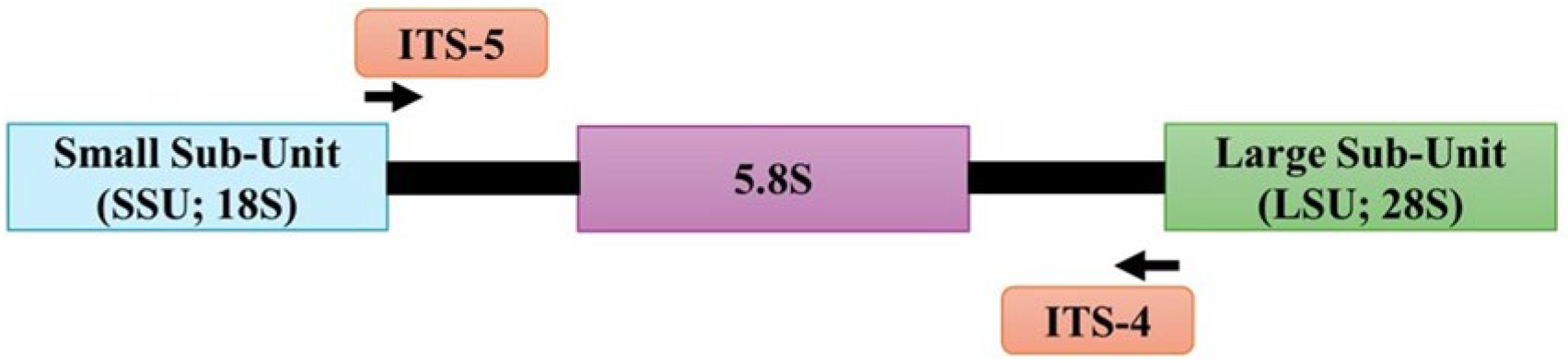
Location of internal transcribed spacer (ITS) primers in fungal ribosomal RNA (rRNA).

There are no recent studies in Mississippi describing the fungal pathogens affecting watermelon production and productivity. Information on prevailing diseases can help guide future breeding strategies and disease management practices. The two research questions addressed by this study are: (i) What is the diversity of putative pathogenic fungi in watermelons in Southwest Mississippi? and (ii) What is the phylogenetic relationship of the characterized isolates? The findings of this research serve as the foundational knowledge for further research on pathogenesis and disease management approaches.

## Materials and Methods

### Sample collection and processing

A watermelon research field at the Alcorn Experiment Station, Lorman, Mississippi (31°52’23.1”N; 91°07’47.0”W) was scouted for the presence of diseases during summer 2024. The research plots had 14 seeded watermelon genotypes. Seeds for those genotypes were obtained from the United States Department of Agriculture germplasm repository at Griffin, Georgia [25, 26] and represented three species of watermelon: *Citrullus lanatus, C. mucosospermus*, and *C. amarus*. A check variety (seedless hybrid named ‘Watermelon Ultra Cool Hybrid’ or ‘WUCH’) was also part of the research.

Symptomatic leaves were collected from eight plots, and a symptomatic fruit was collected from one additional plot. The standing crop was in its fruit set to fruit sizing stage (60 days after seeding) at the time of sampling. Each plot was 6.5 feet long and 1.33 feet wide with five plants per plot for each genotype. Each symptomatic sample (leaf or fruit) was harvested using scissors sterilized with 70% ethanol, kept in plastic Ziploc bags, and then placed in a cooler containing ice. At least three leaves were collected from each sampled plot, but only one representative leaf was used for subsequent fungal isolation.

Among the nine collected samples, five samples belonged to *C. lanatus* domesticated watermelon, two samples were from *C. amarus* wild watermelon, and two samples were from *C. mucosospermus* wild watermelon. The four *C. lanatus* genotypes from which five samples were collected were plant introduction (PI) 635598 (Golden Midget), Watermelon Ultra Cool Hybrid (WUCH), PI 674463 (Moon & Stars), and PI 635699 (Charleston Diploid 59-1). The two *C. amarus* genotypes from which samples were collected were PI 255136 (Tsamma) and PI 688009 (PI 296341-FR). Similarly, the *C. mucosospermus* genotype, PI 560013 (Egusi), was the source of one leaf and one fruit sample collected from the watermelon field. The samples were arranged in the laboratory and photographed along with a ruler to denote the lesion size (Figure 2).

**Figure 2.**
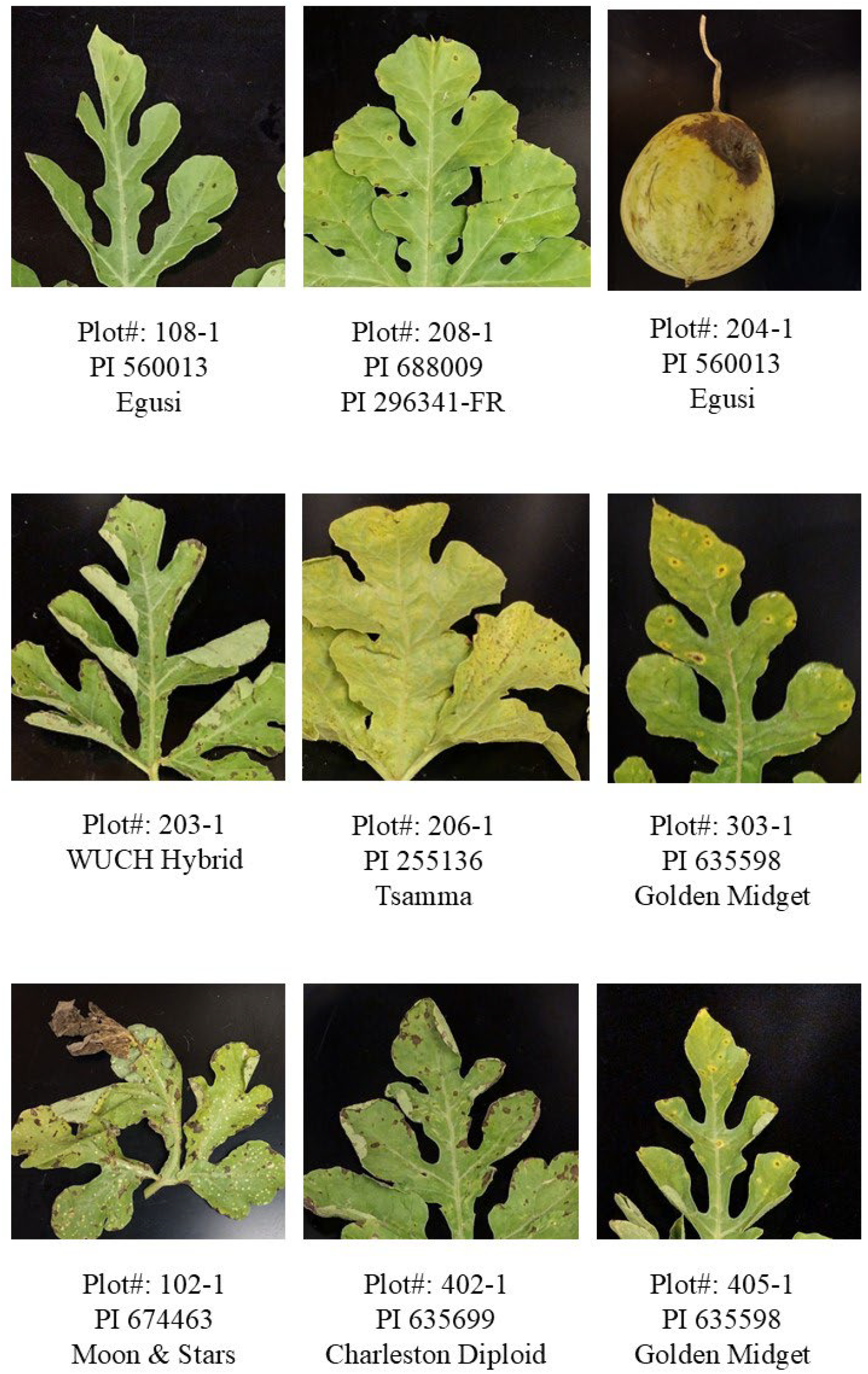
Symptomatic leaf and fruit samples collected from the watermelon research field.

Small sections (1-2 mm^2^) of the symptomatic leaf and fruit samples were excised using a sterile scalpel inside a biosafety cabinet. The cut sections included the margin between the healthy tissue and the progressing symptomatic tissue, avoiding the core necrotic growth. Surface sterilization of the samples was done using 1% sodium hypochlorite for 1 min, followed by two rinses with sterile water for 30 s, and eventually dried on a sterile filter paper. The sterilized, dried leaf or fruit samples were placed on sterile, full-strength potato dextrose agar (PDA) media. Four sterilized samples (one in each quadrant of the media) were placed per PDA plate. The PDA plates were incubated at room temperature (24 °C), and fungal growth was monitored for the next two days. Once the fungal growth was observed on the PDA plates, the growing mycelia were excised using a sterile scalpel, placed in fresh PDA plates, and incubated at room temperature (24 °C).

### Preliminary observations, subculturing, and storage

Fungal growth was monitored on the PDA plates for 1-2 days. Subculturing of the fungus was done by cutting fungal sections from each original PDA plate using a sterile scalpel and subsequently transferring those sections to fresh PDA plates. At least two rounds of subculturing were done until a pure culture of each fungal morphotype was observed. Altogether, nine unique fungal morphotypes were obtained from these subculturing. The pure fungal cultures were cryo-preserved by placing small mycelial sections (fungal mass) in sterile 30% glycerol and kept in - 80 °C freezer for long-term storage.

### Fungal DNA extraction and quantification

Cryopreserved fungal sections were cultured on fresh PDA plates for DNA extraction. Fungal DNA extraction was done using DNeasy Plant Pro Kit (Qiagen, Hilden, Germany) using the manufacturer’s instructions. For maceration of fungal tissue, liquid nitrogen was added to the fungal mycelia placed on a sterile mortar and then ground using a sterile pestle. A minimum of ∼ 100 mg of fungal tissue was taken for DNA extraction. The extracted fungal DNA was assessed for quantity (concentration) and quality (260/280 and 260/230 ratios) using a NanoDrop machine (Thermo Scientific, Delaware, USA).

### Polymerase chain reaction, purification, and sequencing

The fungal internal transcribed spacer (ITS)-specific markers (ITS-5: 5’ GGA AGT AAA AGT CGT AAC AAG G 3’ and ITS-4: 5’ TCC TCC GCT TAT TGA TAT GC 3’) were used for the polymerase chain reaction (PCR) using fungal DNA as the template. The components of the PCR (1X) included molecular biology grade water (8.5 µL), GoTaq Master Mix (12.5 µL), ITS-5 primer (1 µL), ITS-4 primer (1 µL), and fungal DNA (2 µL), totaling 25 µL reaction for each fungal sample. A blank sample (or negative control without the DNA) was also used in the PCR to check the presence of any spurious bands. The PCR was conducted in a T100 Thermal Cycler (Bio-Rad, California, USA) using the program: one cycle (initial denaturation at 95 °C for 2 min), followed by 30 cycles (denaturation at 95 °C for 30 s, primer annealing at 58 °C for 30 s, and extension at 72 °C for 2 min), one cycle of final extension at 72 °C for 10 min, and held at 12 °C. Post PCR, 5 µL of the PCR product for each sample was loaded in a 1% agarose gel stained with 10 µL of SYBR™ Safe DNA gel stain (Invitrogen, Thermo Fisher Scientific). Five positive control samples (products obtained from ITS-primers based PCR using mushroom DNA as a template) and a blank (negative control containing PCR master mix and primers but without DNA) were also loaded in the gel. Likewise, a one-kilobase (kb)-plus DNA ladder (New England Biolabs) was used as a marker in the gel. The gel electrophoresis settings were: 90V, 400 milli Amperes (mA) for 40 minutes. The gel was visualized under an ultraviolet (UV)-transilluminator.

Purification of the PCR products was carried out using Monarch PCR & DNA Cleanup Kit (New England Biolabs, Ipswich, MA, USA) using the manufacturer’s instructions. The purified PCR samples were stored in a +4 °C refrigerator until shipping. The quantity and quality of the purified PCR product was checked. Twelve µL of each purified PCR sample was aliquoted into well-labeled, sterile 200 µL PCR strip tubes and sent for sequencing to Plasmidsaurus Inc., (Kentucky, USA). A linear/PCR sequencing approach was conducted for each sample using Oxford Nanopore technology.

### Processing, analysis, and deposition of the sequenced reads

The sequence data obtained was processed using Geneious Prime^®^ version 2025.0.3. Similarly, the trimmed sequences were aligned using the National Center for Biotechnology Information (NCBI) platform. Also, phylogenetic trees for each fungal isolate were constructed using Molecular Evolutionary Genetics Analysis (MEGA) version 11.0.13 [27]. Trimming of sequencing files in ‘.ab1’ format was performed using Geneious Prime^®^ version 2025.0.3 to remove low-quality reads. The ‘Annotate and Predict’ function within the Geneious Prime software was used to trim the reads with an error probability limit set at 0.01 (1% chance of error per base). The trimmed sequences of the fungal isolates were subjected to alignment using the ‘core_nt’ and internal transcribed spacer (ITS) database available online at the NCBI nucleotide Basic Local Alignment Search Tool (BLASTn) (https://blast.ncbi.nlm.nih.gov/Blast.cgi) [28]. Also, the trimmed ITS sequences for all nine fungi were submitted to the NCBI GenBank database (Table 3).

### Construction of phylogenetic trees

Sequence of top matching accessions from the ‘core_nt’ and ‘ITS’ database (Table 3) and sequence from nine strains from this study were initially compiled in a text file. All these sequences were aligned using ‘Align’ function within the MEGA11 software [27] through MUSCLE version 3.8.425 [29]. This was followed by the generation of a maximum-likelihood phylogenetic tree in MEGA11. The settings used were 1000 bootstrap replicates and the default Tamura-Nei model since it considers the substitution rate between nucleotides and can also handle unequal nucleotide frequencies [30]. The branch length in trees depicted the number of substitutions per site. The tree had a scale showing the nucleotide substitutions. The internal nodes of the trees had bootstrap values showing how many times out of 100 instances (bootstrap replicates), the same clade of the tree could be recovered.

## Results

### Observations on fungal growth

Pure cultures of the fungal samples obtained in this study had variable growth rate after placement in PDA for eight days at 24 °C. Sample ‘ASU_S9_203-1’ had the least growth after eight days. Samples ‘ASU_S8_204-1’, ‘ASU_S11_303-1’, and ‘ASU_S14_204-1’ had relatively lesser growth than other isolates. Similarly, diverse pigmentation was observed on the PDA plates, resulting in nine unique fungal morphotypes (Figure 3).

**Figure 3.**
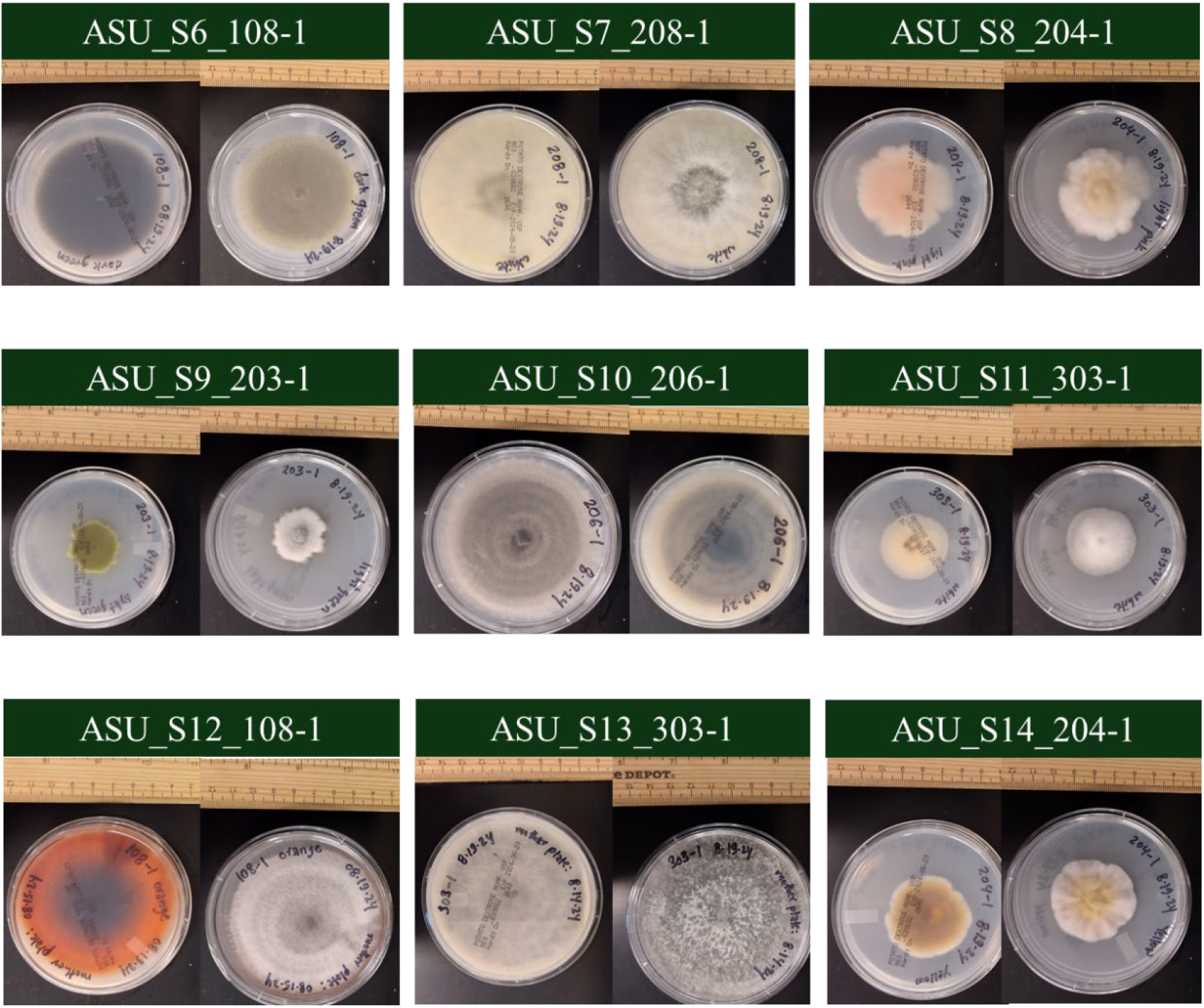
Growth of nine fungal pure cultures eight days after placement on full strength potato dextrose agar (back and front view of the PDA plates for each isolate).

### Quality check of the PCR product

All the positive controls and samples in this study (ASU_S6 to ASU_S14) amplified at the expected ∼600 bp length and showed clear bands indicating successful PCR. As expected, the negative control did not have any band (Figure 4).

**Figure 4.**
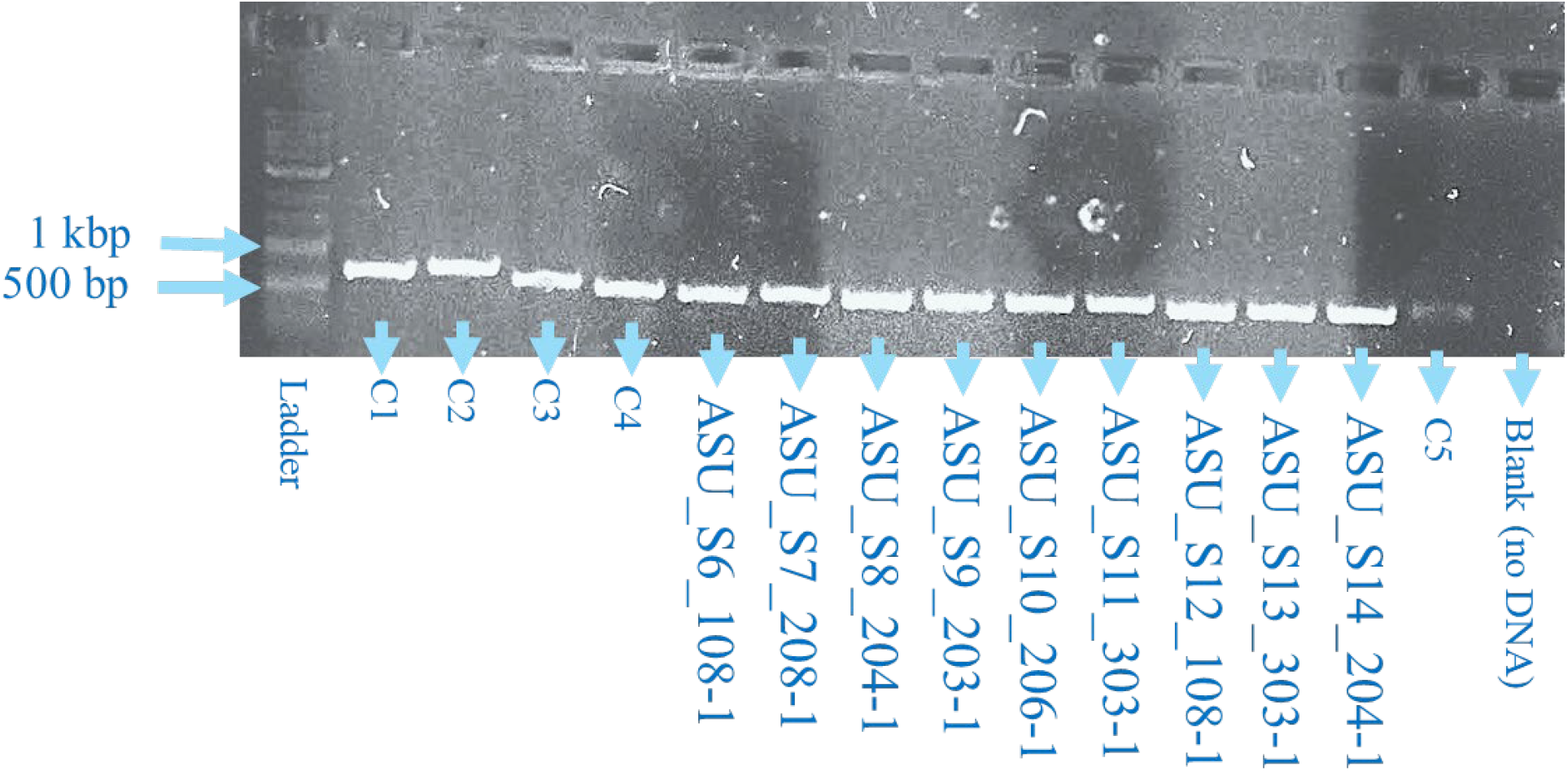
Gel electrophoresis of fungal internal transcribed spacer (ITS) PCR products (ASU_S6 to ASU_S14: fungal isolates from this study; C1-C5: ITS PCR product using mushroom DNA template; Blank: negative control; bp: base pairs; kbp: kilobase pairs).

### Quality check of the purified PCR product

All the purified PCR products showed a concentration between 48 to 64 ng/µL. The absorbance 260/280 ratios were between 1.8 to 2.0. Only one sample ‘ASU_S6_108-1’ had the 260/280 absorbance value slightly above 2. The 260/230 ratios were found to be around the desired value of 2.0 to 2.2. Sample ‘ASU_S7_208-1’ had the least 260/230 value, whereas the sample ‘ASU_S12_108-1’ had the highest 260/230 value (Table 1).

**Table 1.** Quantification of purified PCR products before sequencing.

| Sample ID | Concentration (ng/μL) | 260/280 | 260/230 |
| --- | --- | --- | --- |
| ASU_S6_108-1 | 52.8 | 2.08 | 2.31 |
| ASU_S7_208-1 | 49.0 | 1.96 | 1.88 |
| ASU_S8_204-1 | 60.2 | 1.99 | 2.17 |
| ASU_S9_203-1 | 53.8 | 1.98 | 2.20 |
| ASU_S10_206-1 | 57.2 | 1.95 | 2.33 |
| ASU_S11_303-1 | 54.4 | 1.94 | 2.06 |
| ASU_S12_108-1 | 57.6 | 1.96 | 2.40 |
| ASU_S13_303-1 | 48.7 | 1.98 | 2.11 |
| ASU_S14_204-1 | 63.8 | 1.97 | 2.17 |

### Sequencing summary

All the samples submitted for sequencing successfully passed the Oxford Nanopore sequencing platform. The sequencing results contained several file formats including ‘.ab1’ (chromatogram) file. The trimmed sequences showed that the trimming ranged from a single base (∼ 0.17%) in ASU_S7_208-1 and ASU_S11_303-1 to nine bases (1.66%) in ASU_S6_108-1 (Table 2).

**Table 2.** Length of the sequenced fungal internal transcribed spacer (ITS) reads before and after trimming (bp = base pairs).

| Sample ID | Pre-Trimming<br>Sequence Length (bp) | Post-Trimming<br>Sequence Length (bp) | % Trimmed |
| --- | --- | --- | --- |
| ASU_S6_108-1 | 542 | 533 | 1.66 |
| ASU_S7_208-1 | 564 | 563 | 0.17 |
| ASU_S8_204-1 | 533 | 529 | 0.75 |
| ASU_S9_203-1 | 522 | 519 | 0.57 |
| ASU_S10_206-1 | 576 | 567 | 1.56 |
| ASU_S11_303-1 | 581 | 580 | 0.17 |
| ASU_S12_108-1 | 513 | 512 | 0.19 |
| ASU_S13_303-1 | 534 | 528 | 1.12 |
| ASU_S14_204-1 | 544 | 543 | 0.18 |

**Table 3.**
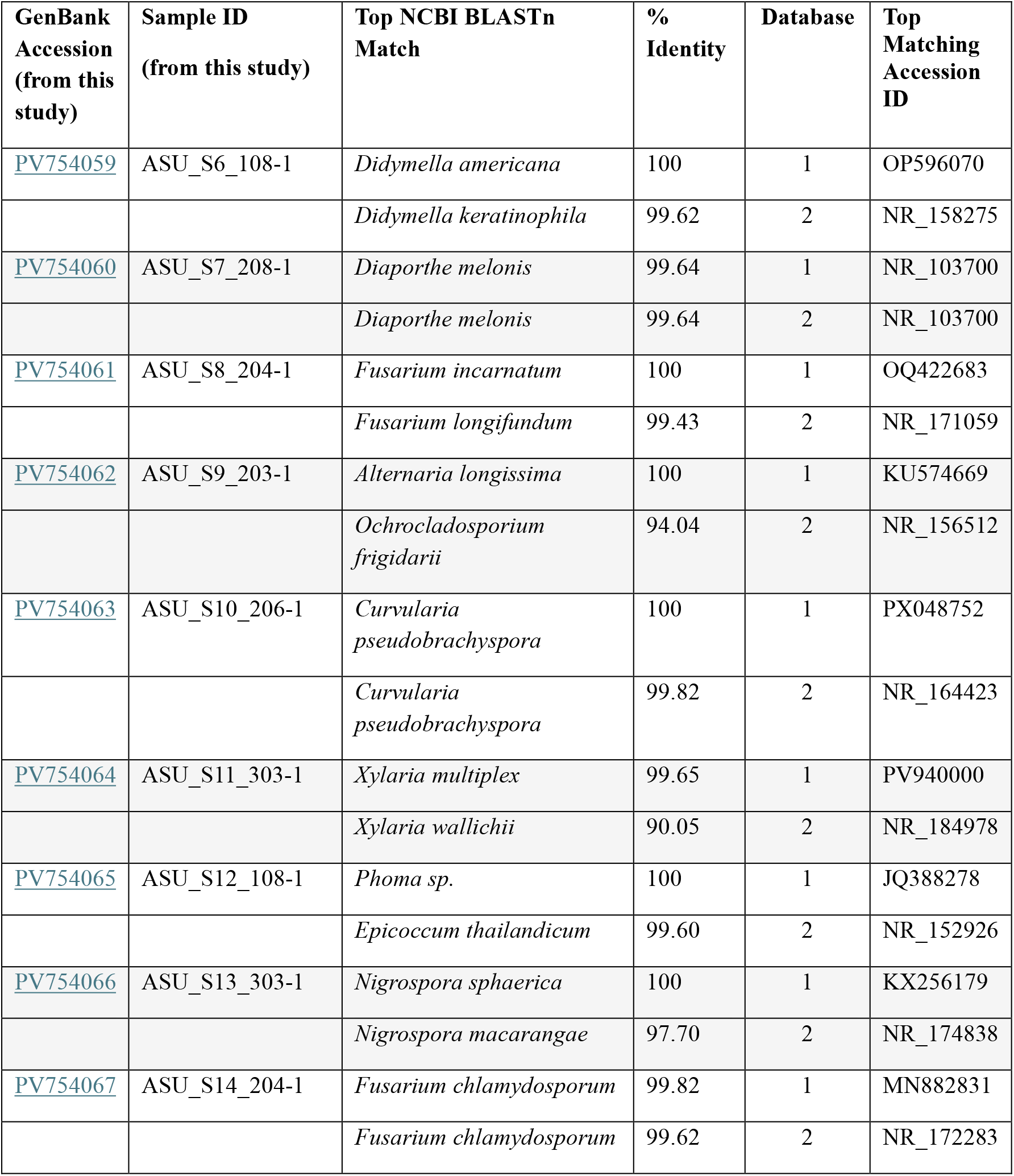
Alignment of the trimmed fungal internal transcribed spacer (ITS) sequences with NCBI BLASTn ‘core_nt’ (1) and ‘ITS’ (2) database. All expect (E) values for the top matches were ‘0’.

### Sequence alignment results

The NCBI BLASTn analysis using ‘core_nt’ and ‘nr/nt’ database was able to resolve the genus and tentative species of the fungal isolates (Table 3). Both the databases showed similar genera for a single sequence except for ASU_S9_203-1 and ASU_S12_108-1. This is common because the ‘core_nt’ database comprises both curated and non-curated sequences, whereas the ITS database strictly contains only the curated sequences and has significantly lesser number of sequences in its repository than the former database. All nine fungal samples in this study were ascomycetes at the phylum-level. Four fungal isolates (ASU_S6_108-1, ASU_S9_203-1, ASU_S10_206-1, and ASU_S12_108-1) belonged to the order Pleosporales and two isolates (ASU_S8_204-1 and ASU_S14_204-1) belonged to the order Hypocreales. Fungal isolates ‘ASU_S7_208-1’, ‘ASU_S11_303-1’, and ‘ASU S13_303-1’ belonged to the order Diaporthales, Xylariales, and Trichosphaeriales, respectively.

### Evolutionary relationship of fungal strains

The maximum-likelihood based phylogenetic tree showed that all the fungal strains grouped in the clade of fungal accessions obtained from the ‘core_nt’ results of BLASTn (Figure 5).

**Figure 5.**
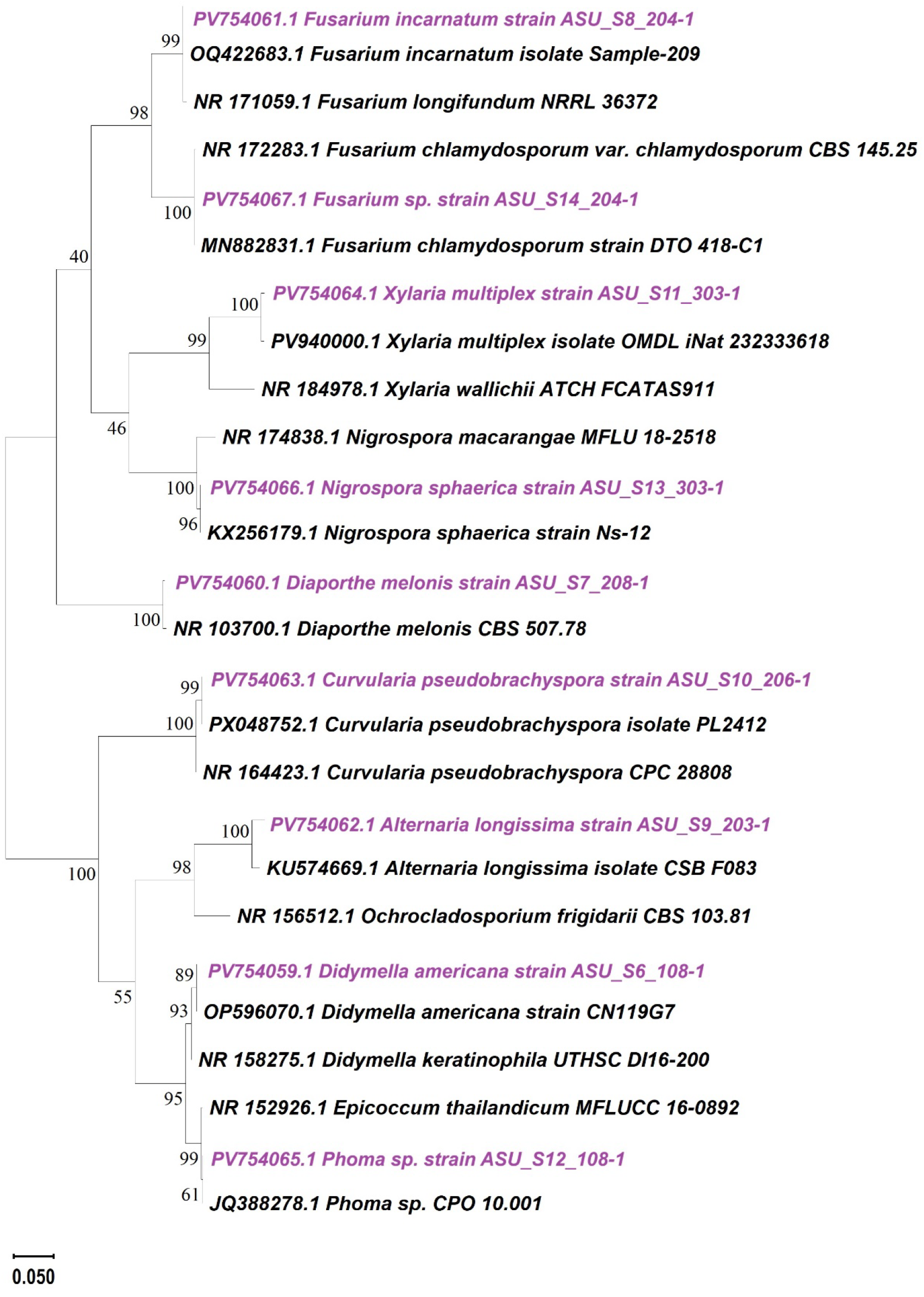
Phylogenetic tree for nine fungal strains from this study (*highlighted with purple*) and their top BLASTn ‘core_nt’ and ‘ITS’ matches constructed in MEGA11 software using maximum-likelihood method. The internal nodes depict the bootstrap values in percentage. A scale bar with numbers shows nucleotide substitutions per site.

## Discussion

Nine unique potentially pathogenic fungal morphotypes were identified in this study. The identification of these putative pathogens was confirmed through ITS-based PCR and sequencing. The first potential pathogen identified in our study was *Didymella americana*. In China, *D. americana* was reported on *Cassia nomame*, where it affected more than 80% plant during the mid and later stages of plant growth [31]. Additionally, this pathogen was identified in the United States on corn roots and soybean pods [32], baby lima bean [33], and table beet [34]. In India, *D. americana* was reported to cause leaf blight in lily [35]. This pathogen has also been reported in wheat [36]. The second potential pathogen identified in our study was *Diaporthe melonis. D. melonis* has been reported to cause fruit rot and stem canker in cucumbers [21] and pod/stem blight in soybean [37]. The host range of *D. melonis* includes *Annona squamosa* [38], *Carapa guianesis* [39], cantaloupes (*Cucumis melo*) [40], and sorghum [41]. Another pathogen identified in our study were two species of *Fusarium*. Pathogenic *Fusarium* species have been widely reported in diverse crops. These include *Fusarium* sp. in corn [42] and tomato [43], *F. solani* f. sp. *cucurbitae* in zucchini [44], *F. solani* f. sp. *cucurbitae* race 1 in pumpkin [45], *F. solani* in muskmelon [46], *F. citrullicola* and *F. melonis* in watermelon and muskmelon respectively [47], and *F. chlamydosporum* in hot pepper, eggplant [48], and watermelon [49].

This study identified *Alternaria longissima* as one of the pathogenic fungal species of watermelon. *A. longissima* has been reported to cause stem and leaf blight on sunflowers and leaf spot and stem necrosis on *Sesamum* [50]. We also identified *Curvularia pseudobrachyspora* as one of the potential pathogenic fungi. *C. pseudobrachyspora* has been reported to cause leaf spot disease on cucurbits [20], coconut seedlings [51], areca palm [52] and banana [53], and bulb rot on lilies [54]. Another fungal species identified in this study was *Xylaria multiplex. X. multiplex* is usually considered saprobic and has been associated with wood decay resembling soft rot [55-57]. We also identified *Phoma* sp. in this research. Different species of *Phoma* have been reported in several cucurbits, for instance black rot caused by *P. cucurbitacearum* on bitter melon [58], pink root of muskmelons and watermelons caused by *P. terrestris* [59], and stem blight of cucumber caused by *P. exigua* [60]. We also obtained *Nigrospora sphaerica* as a potential pathogenic fungus on watermelon. This pathogen has a broad host range [61] and was recently reported to cause leaf spots on watermelon [62].

## Conclusions

Overall, this study provides new insights into watermelon-associated fungi in Southwest Mississippi. Nine unique putative pathogenic fungi associated with watermelons have been identified in this study. Particularly, the genera of *Didymella, Fusarium, Diaporthe, Alternaria, Curvularia*, and *Phoma* could be of further importance and investigation. Recommended future studies include pathogenicity assays to confirm Koch’s postulates and determine host range of each of the fungal isolates. Likewise, a polyphasic approach of identification such as microscopy (histopathology) can be utilized. Further, the whole genome sequencing on these fungal isolates would enable us to develop a deeper understanding of the fungal effectors and genes. Genetic studies to understand the mode of inheritance of resistance can be carried out. The genetic basis of underlying disease resistance can also be delineated. Management strategies to combat emerging pathogens and diseases on watermelons/related cucurbits must be developed to ensure high levels of yield and quality of the produce.

## Acknowledgments

The authors would like to acknowledge the support of the College of Agriculture and Applied Sciences (CAAS) at Alcorn State University. We would like to thank the United States Department of Agriculture (USDA) Agricultural Research Service (ARS), Griffin, Georgia for providing the watermelon seeds for research. We highly appreciate Hardy Diagnostics (Santa Maria, CA) for providing PDA media plates used in initial fungal isolation. We also thank Ms. Amisha Bhatt for her technical help in constructing the phylogenetic tree generated in this study.

## Author Contributions

***Conceptualization***, B.P.B.; ***methodology***, B.P.B.; ***software***, Z.B., P.B., G.T., L.B., and B.P.B.; ***validation***, B.P.B.; ***formal analysis***, Z.B., P.B., and B.P.B.; ***investigation***, B.P.B.; ***resources***, F.M. and B.P.B.; ***data curation***, B.P.B.; ***writing—original draft preparation***, Z.B., P.B., G.T., L.B., and B.P.B.; ***writing—review and editing***, F.M. and B.P.B.; ***visualization***, B.P.B.; ***supervision***, B.P.B.; ***project administration***, B.P.B.; ***funding acquisition***, F.M. and B.P.B. All authors have read and agreed to the published version of the manuscript.

## Data Availability Statement

Data supporting the findings is available within the manuscript.

